# Comparative analysis of codon usage patterns in SARS-CoV-2, its mutants and other respiratory viruses

**DOI:** 10.1101/2021.03.03.433699

**Authors:** Neetu Tyagi, Rahila Sardar, Dinesh Gupta

## Abstract

The Coronavirus disease 2019 (COVID-19) outbreak caused by Severe Acute Respiratory Syndrome Coronavirus 2 virus (SARS-CoV-2) poses a worldwide human health crisis, causing respiratory illness with a high mortality rate. To investigate the factors governing codon usage bias in all the respiratory viruses, including SARS-CoV-2 isolates from different geographical locations (~62K), including two recently emerging strains from the United Kingdom (UK), i.e., VUI202012/01 and South Africa (SA), i.e., 501.Y.V2 codon usage bias (CUBs) analysis was performed. The analysis includes RSCU analysis, GC content calculation, ENC analysis, dinucleotide frequency and neutrality plot analysis. We were motivated to conduct the study to fulfil two primary aims: first, to identify the difference in codon usage bias amongst all SARS-CoV-2 genomes and, secondly, to compare their CUBs properties with other respiratory viruses. A biased nucleotide composition was found as most of the highly preferred codons were A/U-ending in all the respiratory viruses studied here. Compared with the human host, the RSCU analysis led to the identification of 11 over-represented codons and 9 under-represented codons in SARS-CoV-2 genomes. Correlation analysis of ENC and GC3s revealed that mutational pressure is the leading force determining the CUBs. The present study results yield a better understanding of codon usage preferences for SARS-CoV-2 genomes and discover the possible evolutionary determinants responsible for the biases found among the respiratory viruses, thus unveils a unique feature of the SARS-CoV-2 evolution and adaptation. To the best of our knowledge, this is the first attempt at comparative CUBs analysis on the worldwide genomes of SARS-CoV-2, including novel emerged strains and other respiratory viruses.

## Introduction

Coronavirus is a globally threatening virus that belongs to the family coronaviridae. The virus family consists of the largest enveloped single-stranded RNA viruses with a genome size ranging from 26-31 kilobases [1], divided into four genera- alpha, beta, gamma and delta coronavirus [2]. The largest genera in coronaviridae is beta-coronavirus that includes HCoV-OC43, HCoV-HKU, SARS-CoV and MERS-CoV [3][4]. The recent outbreak of coronavirus disease 2019 (COVID-19) caused by the Severe Acute Respiratory Syndrome Coronavirus two (SARS-CoV-2) has become a global concern leading to millions of deaths worldwide [5][6][7][8][9][10]. These viruses infect a wide range of hosts from avian to mammalian species, including human, cats, dogs, cows, bats, civets, camels and cause various types of respiratory infections [11][12][13][14]. However, there are several other viruses already known to primarily target the human respiratory system, which includes Respiratory syncytial virus (RSV), Influenza A virus (H1N1 and H3N2 strains), parainfluenza, rhinovirus (RV), adenovirus, and enterovirus, etc. [15]. Although the transmission mode for SARS-CoV-2 is still not clear, all the respiratory viruses share a similar primary mode of transmission (i.e., respiratory droplets or secretions) among the infected individuals. The spread of infection is more rapid in SARS-CoV-2 than SARS-CoV, MERS, RSV and Influenza A viruses. The rapid transmission rate makes the SARS-CoV-2 pandemic a major worldwide concern [16].

Three major clades of SARS-CoV-2 identified from the public database Global Initiative on Sharing Influenza Data (GISAID) were G (mutation of the spike protein S-D614G), V (mutation of ORF3a coding the protein NS3-G251) and S (mutation ORF8-L84S) clades [17]. Currently, among the SARS-CoV-2 genome sequences, G clade and its offsprings GH (ORF3a:Q57H variant) and GR (Spike D614G and Nucleocapsid RG203KR mutations) accounting about for 74% of the total world sequences. Although research has confirmed the low variability of SARS-CoV-2 genomes [18][19], it is still not clear if the fatality rate and transmission difference in various countries may be the outcome of the virulence of clade’s difference [20]. Therefore to get more insights into the pathogenesis and virulence of SARS-CoV-2, it is crucial to analyse the publically available SARS-CoV-2 genomic sequences coming from different geographical locations[21]. These clades were classified based on different variants found in SARS-CoV-2 genomes. A novel SARS-CoV-2 variant VUI202012/01 emerged in South-East England in November 2020, 56 % more transmissible than the previous SARS-CoV-2 variants. This variant has 17 mutations; the most significant one is in the spike protein, i.e., N501Y mutation as spike protein binds to the human ACE2 receptor; it becomes more infectious transmissible between people. Although the disease severity associated with this variant is not clearly understood.

Nevertheless, higher transmissibility leads to increased disease incidence [22]. Another recently identified SARS-CoV-2 variant in South Africa described a new lineage of SARS-CoV-2 501Y.V2, with eight different non-synonymous spike protein mutations, including the K417N, E484K, and N501Y at the key site of the receptor-binding domain. The South Africa variant is also rapidly spreading, although the complete significance is yet to be determined. It is important to check whether these reported mutations in different parts of the virus genome affects codon usage in SARS-CoV-2 [23].

Codon Usage Bias (CUB) is a crucial phenomenon in virus evolution [24]. The genome-wide CUB analysis helps identify selective forces responsible for shaping the codon usage bias and important to related genomic studies [25]. The CUB patterns have been studied recently, especially in the case of different virus genomes [26][27][28]. Many previous studies suggest that several factors are associated with the specific CUB patterns, including nucleotide composition [29], mutational pressure or natural selection [30][31], G+C content [32], protein secondary structure [33], translation process [34], hydrophobicity [35], tRNA abundance [36], environmental temperature [37] and other factors. Relative Synonymous Codon Usage (RSCU), Effective Number of Codons (ENC), neutrality plot, Codon Adaptation Index (CAI), the different selection pressures (mutational or natural selection) and other factors help in the assessment of measuring CUB in different species [38][39][40][41].

A limited number of studies have been published related to CUBs in SARS-CoV-2. For example, Gu et al.’s work on the genus betacoronavirus compared the codon usage pattern for viruses in this genus, using multivariate analysis [42]. Hou performed analysis on the SARS-CoV-2 with human and non-human coronaviruses to determine if different hosts show variable CUBs [43]. Another research was conducted to study the factors influencing codon usage profiles of SARS-CoV-2 in humans and dogs [44]. However, these analyses are performed on a limited number of genome sequences and do not focus on the comparative analysis concerning other known respiratory viruses. We also studied the newly emerged SARS-CoV-2 variants from the UK and South Africa to check if these SARS-CoV-2 mutations are associated with altered codon usage patterns.

The present study aims to understand the patterns and factors that are responsible for governing the CUBs in ~62K genomes distributed across the seven SARS-CoV-2 genomic clades (G, GH, GR, L, O, S and V) and the recently emerged variants (VUI202012/01 and 501Y.V2 from UK and SA). We performed the systematic identification and comparison of CUBs in SARS-CoV-2, host *(H. sapiens)* and other respiratory viruses, i.e., SARS-CoV, MERS-CoV, RSV, H1N1 and H3N2.

## Material and Methods

### Retrieval of genomic sequences

The complete and high coverage FASTA formatted genomic sequences of SARS-CoV-2 clades (G, GH, GR, L, O, S and V), the recently discovered variants VUI202012/01 (UK) and 501Y.V2 (SA), retrieved from GISAID [45], have been used for the study. The complete genome sequences for different respiratory viruses including SARS-CoV-2 (NC_045512.2), SARS-CoV (NC_004718.3), Respiratory Syncytial Virus (RSV, NC_001803), Middle East Respiratory Syndrome coronavirus (MERS-CoV, NC_019843.3), Influenza A virus most common strains (H1N1 and H3N2) and of the host (*H. Sapiens*) were retrieved from the National Centre for Biotechnology virus resources (NCBI) (https://www.ncbi.nlm.nih.gov/labs/virus/).

### Nucleotide Composition Analysis

The nucleotide composition and other codon usage indices for all the respiratory virus genomic sequences were calculated using CodonW. The computed properties include the overall GC content; Effective number of codons (ENC); Codon Adaptation Index (CAI); frequency of each nucleotide at the third position of the synonymous codons (A3s, T3s, G3s, C3s); G+C content at 1^st^, 2^nd^ and 3^rd^ codon positions GC1, GC2, GC3, respectively. The value of GC12 (mean GC content at 1^st^ and 2^nd^ position of the codon) and GC3 (GC content at the 3^rd^ position of the codon) of different clades, along with all respiratory viruses, were calculated by using the EMBOSS CUSP program (http://emboss.sourceforge.net). The CAI values were determined using DAMBE 5.0 [46].

### Phylogenetic analysis

A phylogenetic tree was constructed of complete genome sequences using MEGA 6.0 software [47] using the Neighbor-joining (NJ) algorithm and Kimura-2 parameter model on 1000 bootstrap replicates.

### RSCU analysis and heatmap generation

To investigate the factors affecting the synonymous codon usage bias, RSCU values were calculated using CodonW software (http://codonw.sourceforge.net/). The RSCU values were calculated using the formula:

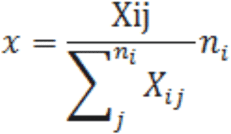

X_ij_ represents the number of codons for the amino acid and ni represents the degenerate number of a specific synonymous codon, ranging from 1 to 61.

High RSCU is the ratio of observed to the expected value for a given amino acid and its value is not affected by the length of the sequence or amino acid frequency [34]. A higher RSCU value (RSCU>1) indicates positive codon bias and is considered as a preferred codon, whereas the lower RSCU value (RSCU<1) represents the negative codon bias termed as under-preferred codons. The RSCU values across all respiratory viruses with respect to host (*H. sapiens*) were compared and visualised with a heatmap in R.

### ENC analysis

To further investigate the synonymous codon usage pattern, the ENC-plot was generated by plotted the ENC values against the GC3 values. The ENC is used to measure the deviation from the random codon usage pattern; its value ranges from 20-61. A lower ENC value (<35) corresponds to a strong codon usage bias, whereas higher ENC values (>35) represent low codon bias [48]. The standard ENC values were calculated using the formula,

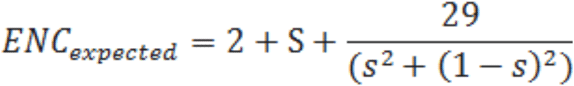

S represents the given GC3s value.

### Neutrality plot analysis

Codon usage disparity is governed by mainly two important factors, mutation pressure and natural selection. In neutrality plot analysis, the main factors affecting the CUBs were determined by taking the mean GC content at 1^st^ and 2^nd^ position (GC12 x-axis) and plotted that against GC content at 3^rd^ position of the codon (GC3 y-axis) values, calculated by CodonW. Plotting of GC12 values against GC3s helps analyse the correlation between the base compositions of all three different codon sites, thus determining the main factor responsible for the codon usage bias. The regression line’s slope indicates the effect of mutational pressure [49].

### Dinucleotide frequency analysis

The dinucleotide frequency analysis is another way of establishing the relation with the codon usage bias, calculated using DAMBE. The average relative abundance value for each dinucleotide was determined by the odds ratio, defined as the ratio of observed and expected dinucleotide frequencies. The odds ratio value >1.23 was considered over-represented, whereas the value <0.78 as under-represented [50].

## Results

We collected 61,962 SARS-CoV-2 whole-genome sequences submitted to the GISAID until April 2020, consisting of 13280, 15506, 20779, 2660, 2615, 4183, and 2939 sequences of G, GH, GR, L, O, S and V clades, respectively. We also collected the recently reported SARS-CoV-2 variant sequences for VUI202012/01(n= 528) and 501Y.V2 (n= 184) from the UK and SA. As expected, the phylogenetic tree indicates that the newly SARS-CoV-2 was evolutionarily closer to the SARS-CoV and MERS viruses. The phylogenetic tree also reveals the SARS-CoV-2 genome to be relatively distant from the influenza A virus strains (H1N1 and H3N2) clustered with the Respiratory Syncytial Virus (RSV) forming a separate clade (**Figure 1a**).

**Figure 1.**
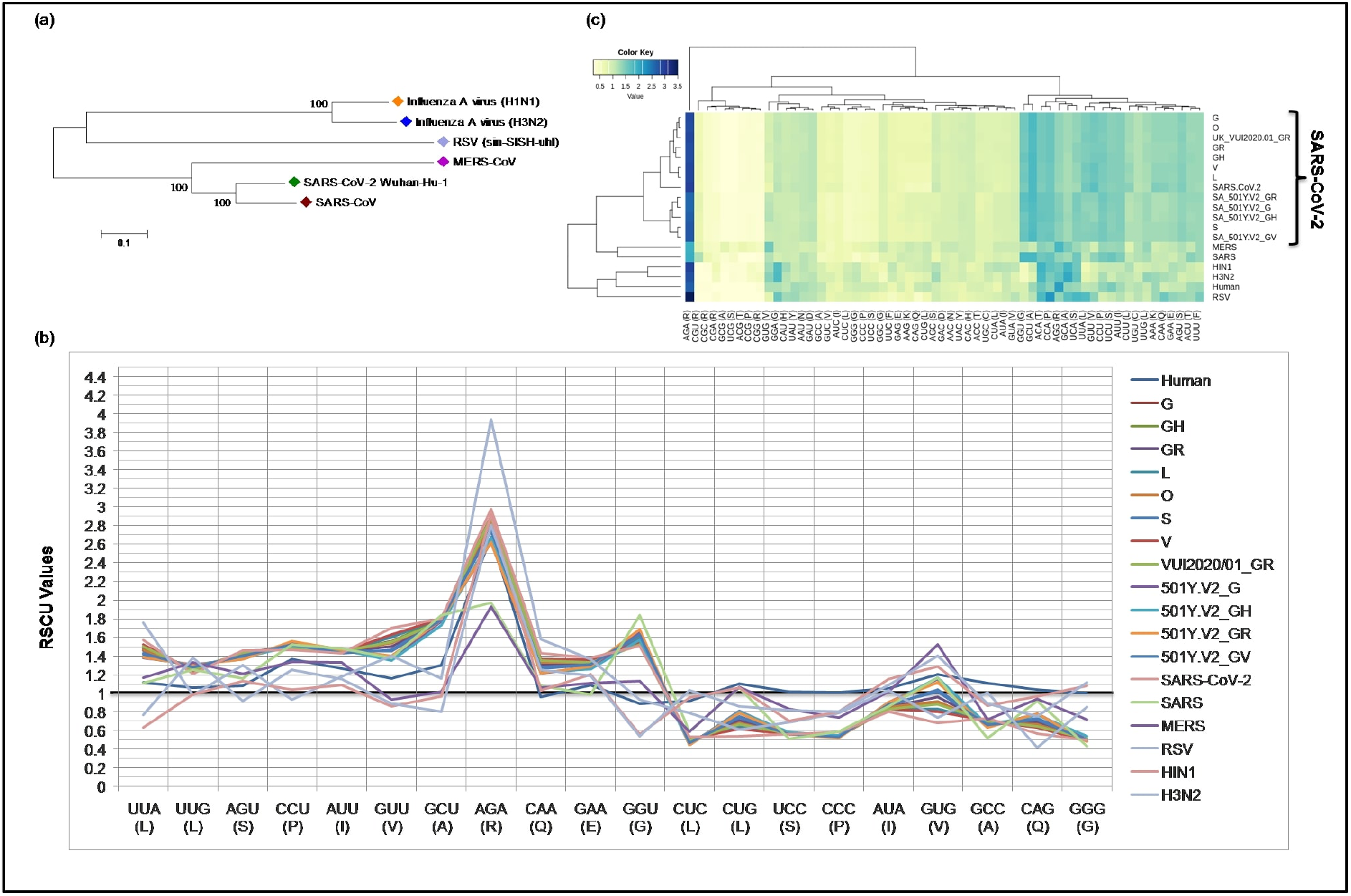
RSCU analysis of the ~62k SARS-CoV-2 genomes and other respiratory viruses. (a) Neighbour-joining phylogenetic tree for all the respiratory viruses included in this study, namely, SARS-CoV-2, SARS-CoV, MERS-CoV, RSV, H1N1, and H3N2. (b) The profiles of the RSCU values of the over and under-preferred codons (RSCU difference >=0.20 between human and SARS-CoV-2 genomes), the bold horizontal line showing the categorisation of RSCU values >1 (Preferred codons), and below 1 (under-represented). (c) Heatmap of the RSCU values for all respiratory viruses genomic sequences. The highly preferred codons are highlighted in blue, the preferred ones highlighted in green and the non-preferred or under-represented in yellow.

### Nucleotide composition and codon usage indices

We calculated the nucleotide composition for all the respiratory viruses genomes studied here. The nucleotide composition for SARS-CoV-2, SARS, and MERS found to be similar and in the order U>A>G>C. Whereas RSV follows the order A>U>G>C, and H1N1 and H3N2 followed the order A>G>U>C. We found that the nucleotides at the 3^rd^ position of the codon follow the trend U_3_>A_3_>C_3_>G_3_ for all viruses, except SARS-CoV and MERS, for which the trend is U_3_>A_3_>G_3_>C_3_ (**see S.Table 1**). The average GC content for all the respiratory viruses is 0.40, with a standard deviation of 0.02. The CAI for all respiratory viruses ranges from 0.68-0.72. The higher CAI value indicates a better adaptation of the virus in the human host.

### Codon usage patterns among SARS-CoV-2 clades and other respiratory viruses

The CUB exists in many RNA viral genomes, generally determined by mutation and selection pressure. RSCU analysis was performed to investigate the codon usage bias variation in all the studied virus genomes. With respect to the *H. sapiens* host, we found 11 over-represented or preferred codons by SARS-CoV-2 clades, i.e., UUA, UUG, AGU, CCU, AUU, GUU, GCU, AGA, CAA, GAA and GGU. Out of these, 6 are ending with U, 4 with A and 1 with G. The 9 under-represented or randomly used codons were CUC, CUG, UCC, CCC, AUA, GUG, GCC, CAG and GGG comprising of 4-G ending, 4-C ending, and 1-A ending (**see S.Table 2**). From the identified codons, most of them are previously reported as preferred codons [51]. From the analysis, we found that all the viruses studied here are highly biased towards A/U-ending codons. In contrast, the under-represented codons were found to be C/G ending, as reported earlier in the recent studies conducted on SARS-CoV-2 genomes [44]. The over and under-represented codons RSCU values concerning host were plotted in a line plot (**Figure 1b**). The RSCU ranges from 0.12 for Alanine to 3.94 for Arginine among the studied genomes. The heatmap and the associated clustering of the over and under-represented codons are shown in **Figure 1c**. All the clades of SARS-CoV-2 genomes along with SARS and MERS clustered together, forming a separate clade.

Among the respiratory viruses, it has been observed that there is no significant CUB pattern observed among the clades. Moreover, a significantly different pattern was observed in all the SARS-CoV-2 clades with reference to the human genome. For a few codons, the RSV and influenza A virus show a slight variation in CUB, compared with that of the other respiratory viruses, as shown in **S.Table2.** The phylogenetic analysis also confirms the pattern as these three viruses clustered together, forming a separate clade. Summarily, our results suggest that the codon usage pattern is highly similar in SARS-CoV-2 clades, including the newly discovered variants from the UK and SA.

### Mutational pressure, the key driving force of codon usage bias among respiratory viruses

To determine whether mutational pressure or natural selection are the key driving factors affecting the codon bias within these respiratory viruses, the ENC-plot was generated by plotting ENC values against the corresponding GC3s values. The ENC-GC3s plot revealed that among all the respiratory viruses, the mean ENC value is 51.47, with a standard deviation of 2.11, suggesting that the codon bias is relatively low among all the respiratory viruses. All the dots in ENC-plot were located below the standard curve (**Figure 2a**), indicating that the codon bias in all the viruses is majorly due to mutational pressure. The minimum ENC value was observed for RSV (46.89), while the maximum for SARS (53.34).

**Figure 2.**
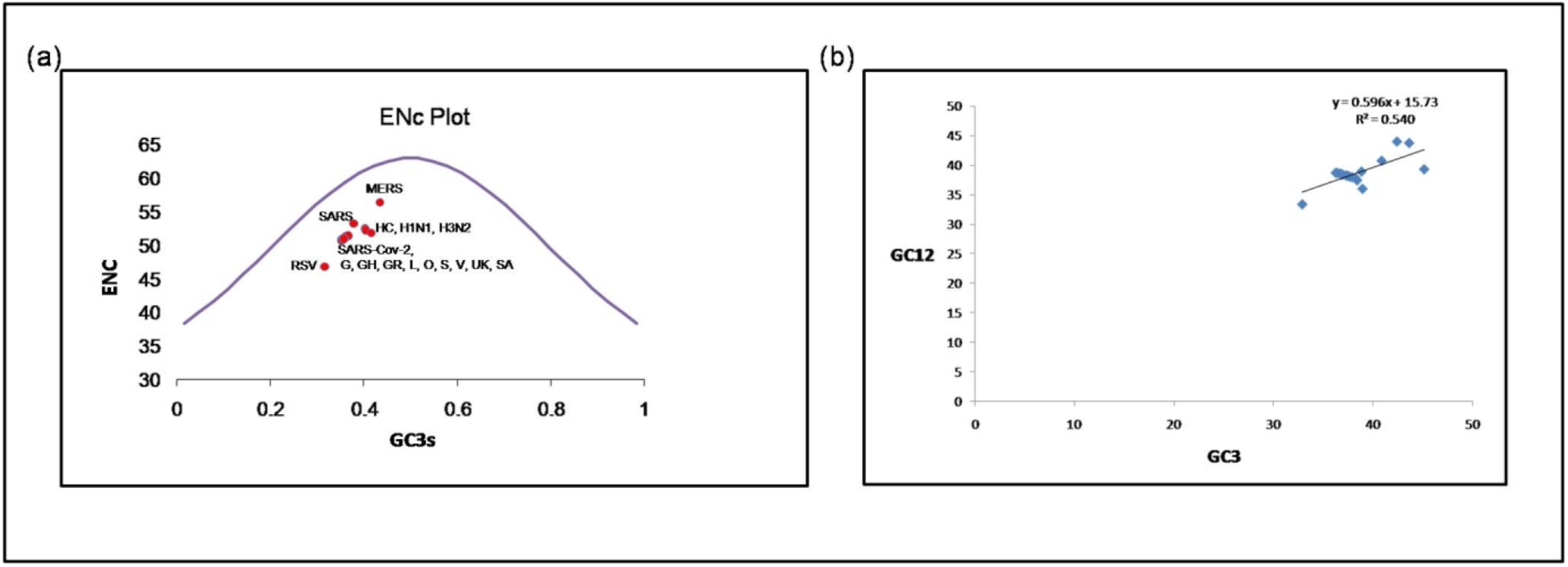
ENC and neutrality plot analysis of all the respiratory viruses in the study. (a) The ENC values plotted against GC3s; the blue curve represents the standard curve in the absence of selection. Red dots show different respiratory viruses included in the study. The ENC and GC3 values obtained for different viruses were found very similar (SD of ± 2.11 and ±0.03, respectively) because of the reason, they clustered. (b) A neutrality plot was generated for respiratory viruses, including the newly emerging SARS-CoV-2, by plotting GC content at 1^st^ and 2^nd^positions of the codon against the GC content at the third position of the codon. Blue dots represent each clade and variants from the UK and SA of the SARS-CoV-2 virus and other respiratory viruses in the plot.

### Neutrality plot analysis

The neutrality plot (**Figure 2b**) indicates a good correlation between the GC12 and GC3 values belonging to different clades of the SARS-CoV-2 virus (r^2^= 0.54) and other viruses. The regression line’s slope and intercept were calculated to be 0.596 and 40.4, suggesting 60% and 40% contribution by mutational pressure and natural selection, respectively.

### Influence of dinucleotide frequency in determining the codon usage bias among the respiratory viruses

As dinucleotide usage is another crucial factor in determining the codon usage bias, we computed the relative dinucleotide abundance value of 16 dinucleotides combinations. The most abundant dinucleotides across all the respiratory viruses were UpG, CpA, ApC, CpU and GpC with odds ratios 1.36, 1.26, 1.169, 1.162, 1.073, respectively. CpG (odds ratio: 0.415) was the least abundant dinucleotide observed in all the respiratory genomes (**S.Table3**). The relative abundance of UpG (1.36) and CpA (1.28) dinucleotides were found to be over-represented compared to others **(Figure 3)**. The RSCU values for seven codons containing UpG (UUG(L), CUG(L), AGU(S), GUU(V), GUG(V), GCU(A), and GGU(G)) and four codons with CpA (UAC(Y), CAU(H), CAA(Q), and AAC(N)) were analysed to identify the possible effect of UpG and CpA representations on codon usage bias.

**Figure 3.**
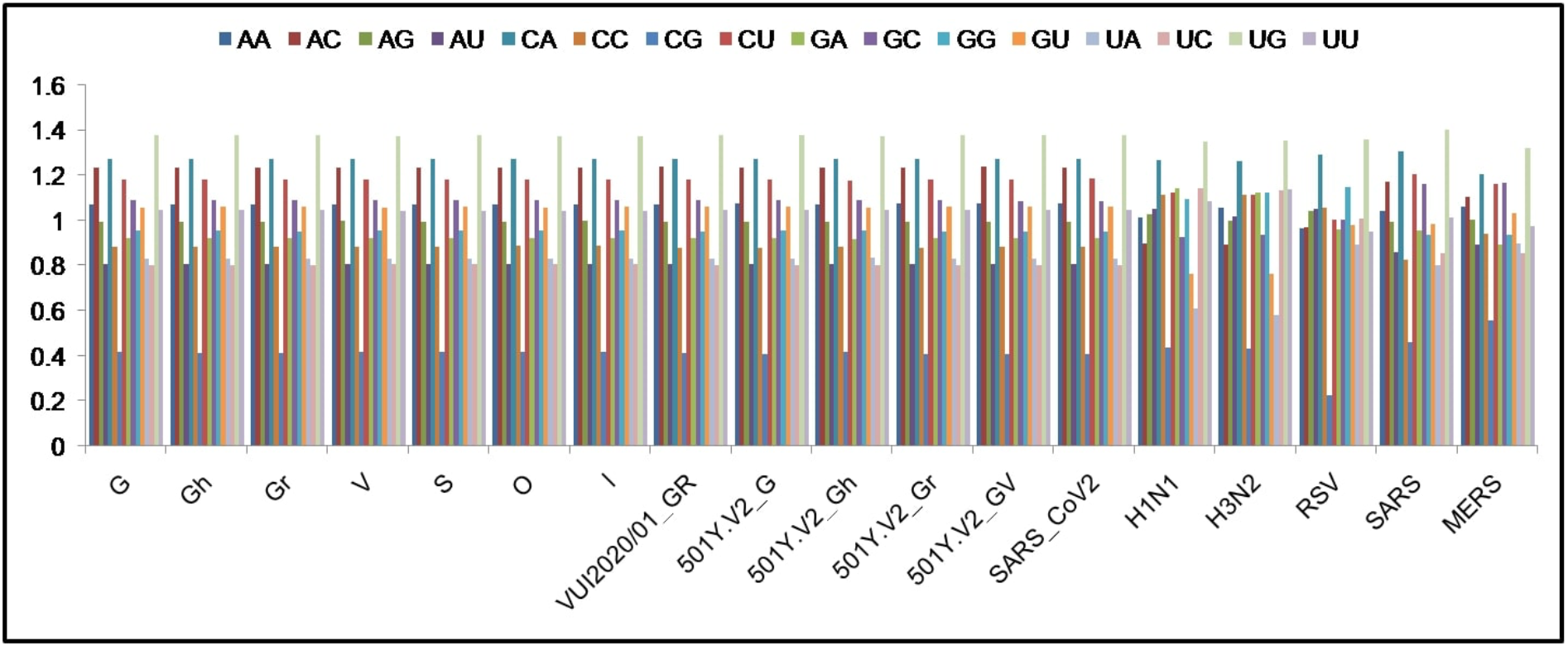
Relative dinucleotide frequencies among all the respiratory viruses.

## Discussion

Codon usage is affected by several factors that include various sequence-based properties such as nucleotide composition, dinucleotide composition, amino acid preferences, host adaption, etc. There is a unique and characteristic ‘signature’ of codon usage bias for each genome. SARS-CoV-2 has a very distinct codon usage pattern as compared to other human beta coronaviruses [52]. We performed this study to investigate the factors governing codon usage bias in all the respiratory viruses, including SARS-CoV-2 isolates from different geographical locations. The analysis includes RSCU analysis, GC content calculation, ENC analysis, dinucleotide frequency and neutrality plot analysis.

From the RSCU analysis for all the viruses studied here, we observed a slight variation in RSCU values amongst all the SARS-CoV-2 genomes. Moreover, compared with the human host, a significant difference in RSCU patterns in all the SARS-CoV-2 genomes were observed. As compared to other viruses included in the study, only RSV and Influenza A show a slight individual variation in the RSCU values. Similar to other RNA viruses, all the respiratory viruses studied here tend to have higher AU content, suggesting the role of mutational pressure in the selection of codon usage [53]. The RSCU analysis for all the respiratory viruses revealed that highly preferred codons (RSCU >1) were A/U ending, while the non-preferred (RSCU <1) codons were G/C ending, which is in agreement with the previous studies [54][51]. Our analysis led to the identification of 11 over-represented codons (UUA (L), UUG (L), AGU (S), CCU (P), AUU (I), GUU (V), GCU (A), AGA (R), CAA (Q), GAA (E) and GGU(G)) and 9 under-represented codons (CUC (L), CUG (L), UCC (S), CCC (P), AUA (I), GUG (V), GCC (A), CAG (Q), and GGG (G)) in all the respiratory virus genomes. Interestingly, Leucine (UUA, UUG), Serine (AGU), Proline (CCU), Isoleucine (AUU), Valine (GUU), Alanine (GCU), Arginine (AGA), Glutamine (CAA), Glutamate (GAA), and Glycine (GGU) codons were preferred by all the ~62K SARS-CoV-2 genomes, while influenza A virus (H1N1, H3N2) preferred Phenylalanine (UUC), Threonine (ACA), Alanine (GCA), and Glycine (CGA) codons. Serine (UCA) and Proline (CCA) codons were preferred by SARS and RSV, whereas RSV virus shares few similar codons of influenza A virus strains, namely, Alanine (GCA) and Glycine (CGA). The codon usage pattern among RSV and influenza A virus is slightly different when compared with other respiratory viruses. In addition to this, SARS and MERS show a variation from the other respiratory viruses for the codons for Arginine (AGA) and Glutamate (GAA).

The nucleotide composition analysis in the study revealed that in SARS-CoV-2, the composition follows the order U>A>G>C. We also observed that U and A nucleotides occurred most frequently at the codon’s third position, confirming a previous finding [43]. All the viruses included in this study show a higher frequency of A and U nucleotides, suggesting that the bias in the genome composition of respiratory viruses also reflects in their codon usage patterns.

To further estimate the degree of codon usage bias among respiratory viruses, ENC values were calculated. In general, the RNA viruses show higher ENC (> 45), suggesting a random codon usage [54]. The mean ENC value of all the viruses studied here is 51.47, with a standard deviation of 2.04, reflecting a relatively low codon usage bias for all the viruses, similar to the previous finding for SARS, MERS, and other human coronaviruses, different from HCoV-NL63 [55]. The lowest ENC value was found for RSV (46.89), indicating the codon bias to be higher in RSV, as compared to that of other viruses studied here.

Our next attempt is to determine the factors involved in shaping the CUBs. Previous studies suggest that codon usage bias is affected by several factors. Out of these factors, two widely accepted factors are mutation pressure and natural selection [55]. Other influencing factors include GC content, hydrophobicity, and GC3 etc. As stated earlier, we found A3 and U3 frequencies to be higher than G3 and C3, suggesting the contribution of mutational force in shaping the codon usage among respiratory viruses. To further investigate the contribution of mutational force, ENC-plot was generated, where the ENC value is plotted against GC3. If the codon usage bias is only affected by the GC3 value, all the points should be exactly on the standard curve. The ENC-GC3 plot has shown that all the points clustered together just below the standard curve, highlighting that G+C compositional play an important role in shaping the codon usage bias in all the respiratory viruses under study. The points below the curve also indicate that some independent factors, other than mutational force, like natural selection, might also play a role in shaping the CUBs in respiratory viruses [51]. Furthermore, to examine the degree of codon bias contributing forces, the neutrality plot was generated. In the plot, GC content at the 3^rd^position of the codon was plotted against mean GC content at the 1^st^ and 2^nd^positions. The regression coefficient (RC) value of RC<0.5, reflecting a more significant role of natural selection, while RC>0.5 suggests the highest impact of mutational pressure. In the neutrality plot, the RC value of 0.54 indicates the role of mutational pressure is slightly higher than natural selection in governing the CUB among the respiratory viruses studied here. Dinucleotide frequency has shown to be another determining factor for the codon usage amongst the RNA viruses [56]. The analysis of dinucleotide frequency for the studied respiratory viruses indicate that amongst the UpG containing codons, five codons (UUG, AGU, GUU, GCU, and GGU) were found to be over-represented and two (CUG, and GUG) under-represented codons. While for dinucleotide CpA: CAA, UAC, and AAC codons were over-represented. The relative abundance of dinucleotides, UpG and CpA in different organisms results from under-represented CpG di-nucleotides [57]. We also observed the odds ratio range to be slightly different for H1N1, H3N2, and RSV compared to that of other respiratory viruses. The low CpG dinucleotide frequency directly correlates with the codon usage bias amongst the respiratory virus to evade the host immunity, most of the viruses maintain low CpG content[58]. These results highlight that compositional constraints are the primary determinant of codon usage bias, since all the respiratory genomes are rich in A/U nucleotides, consistent with earlier findings by Khandia et al. [59]. Our results suggested that dinucleotide frequency also plays a vital role in codon usage bias among respiratory viruses.

## Conclusion

To the best of our knowledge, this is the first attempt at comparative CUBs analysis on worldwide SARS-CoV-2 genomes, including the newly emerged strains and other respiratory viruses. We conclude that there is no significant impact of reported SARS-CoV-2 mutations on the codon usage preferences. Summarily, it was found that in all the respiratory viruses, the codon usage bias is highly similar and relatively low. The study also discloses that mutational pressure and natural selection are the leading forces determining the codon usage bias amongst all the respiratory viruses.

## Conflict of interests

The authors declare that the research was conducted in the absence of any commercial or financial relationships that could be construed as a potential conflict of interest.

## Funding

This work was financially supported by the Department of Biotechnology (DBT), Government of India, grant:BT/PR40151/BTIS/137/5/2021, awarded to DG. Financial support provided by the Indian Council of Medical Research (ICMR), India, to RS as Senior Research Fellowship is duly acknowledged (2019-5850). NT acknowledges GSK fellowship ID 2362 and RCB (Regional Centre for Biotechnology).

## Acknowledgements

All the authors acknowledge ICGEB for providing the necessary infrastructure and facilities for the research.

**S. Table 1.** All codon usage indices calculated for all the respiratory viruses analysed in this study.

**S. Table 2.** RSCU values for all the respiratory viruses analysed in this study.

**S. Table 3.** Dinucleotide frequencies calculated for all respiratory viruses.

## Notes

**Conflict of interest**: Nothing to declare

### Competing Interest Statement

The authors have declared no competing interest.

